# First record of the seagrass *Halophila stipulacea* (Forskkal) Ascherson in the waters of the continental United States (Key Biscayne, Florida)

**DOI:** 10.1101/2024.09.02.610701

**Authors:** Justin E. Campbell, Aarin-Conrad Allen, Danielle C. Sattelberger, Matthew D. White, James Fourqurean

## Abstract

The first record of *Halophila stipulacea* is reported for the continental waters of the United States. In August 2024, a small meadow was identified inside Crandon Marina on Key Biscayne, Florida, USA. Following surveys have revealed that *H. stipulacea* has spread to adjacent areas immediately outside of the marina, often growing either in close proximity to, or interspersed with, the native seagrasses *Thalassia testudinum, Syringodium filiforme*, and *Halodule wrightii*. This serves as an initial report and extends the geographic scope of this introduced species in the Western Atlantic basin.

## Introduction

*Halophila stipulacea*, is a dioecious, euryhaline seagrass native to the waters of the Red Sea, Persian Gulf, and western Indian Ocean. This species of seagrass has drawn attention in recent years as it has expanded its range into nonnative territories. Initially, *H. stipulacea* crossed through opening of the Suez Canal (den Hartog, 1972; Lipkin 1975a, 1975b) and has since spread west throughout the Mediterranean Sea (Van der Velte & den Hartog, 1989; Bianchi & Morri, 2004; Gambi et al., 2009; Sghaier et al., 2011). In recent decades, *H. stipulacea* has begun an invasive spread for the second time, with new reports of the species across the Caribbean.

Here, we report that *H. stipulacea* is now within the continental waters of the United States (Key Biscayne, Florida), thereby extending its northernmost extent in the Western Atlantic basin.

The first report of *H. stipulacea* in the tropical Western Atlantic occurred in 2001-2002 on the island of Grenada (Ruiz & Ballantine, 2004). Ruiz and Ballantine (2004) hypothesized that this spread occurred from a recreational, luxury yacht departing the Mediterranean. Since this initial report, *H. stipulacea* has spread northward to St. Lucia and Dominica (Willette & Ambrose, 2009; Steiner & Willette, 2015). In 2014, *H. stipulacea* was reported off Venezuela (Vera et al., 2014). Since 2017, its spread has been documented throughout the Lesser Antilles, as far north as the U.S. Virgin Islands (Olinger et al., 2017), Puerto Rico and British Virgin Islands (Ruiz et al., 2017). Until recently, reports of *H. stipulacea* have been restricted to the islands southwest of Hispaniola.

*H. stipulacea* is a fast-growing, pioneering seagrass species, that tolerates a wide range of salinities, temperatures, and light levels, and can withstand various disturbance regimes (Willette & Ambrose, 2009; Short et al., 2010). These factors may have contributed to its rapid expansion across the Caribbean (Ruiz & Ballantine, 2004; Olinger et al., 2017; Ruiz et al., 2017; Scheibling et al., 2018). Within its expanded range, some reports suggest that *H. stipulacea* can outcompete and replace native seagrasses like *Halodule wrightii* and *Syringodium filiforme* (Winters et al., 2020), thus fueling concern over its continued spread.

## Methods

*Halophila stipulacea* was first sighted inside Crandon Marina (25.72586 N, 80.15594 W) on Key Biscayne, FL in August 2024. Multiple small patches were initially observed along the shallow (∼2m depth) western margins of the interior seawall. Initial samples were immediately collected by hand and returned to the lab for identification. In late August 2024, the largest meadow inside the marina was surveyed for overall dimensions, seagrass percent cover, and total biomass. Ten small quadrats (0.0625m^2^) were haphazardly tossed within the meadow and seagrass species and percent cover were recorded. Ten seagrass cores (above- and belowground biomass, 0.0625m^2^) were also haphazardly collected by hand for total biomass. These cores were immediately placed in coolers, returned to the lab, rinsed free of sediment and processed for wet mass (g). Haphazard surveys were also conducted outside of the marina, both along the external margin of the marina entrance and the adjacent shallow seagrass flats and mangrove fringe (Fig. 1). Approximate patch dimensions and seagrass community composition were recorded.

**Fig 1.**
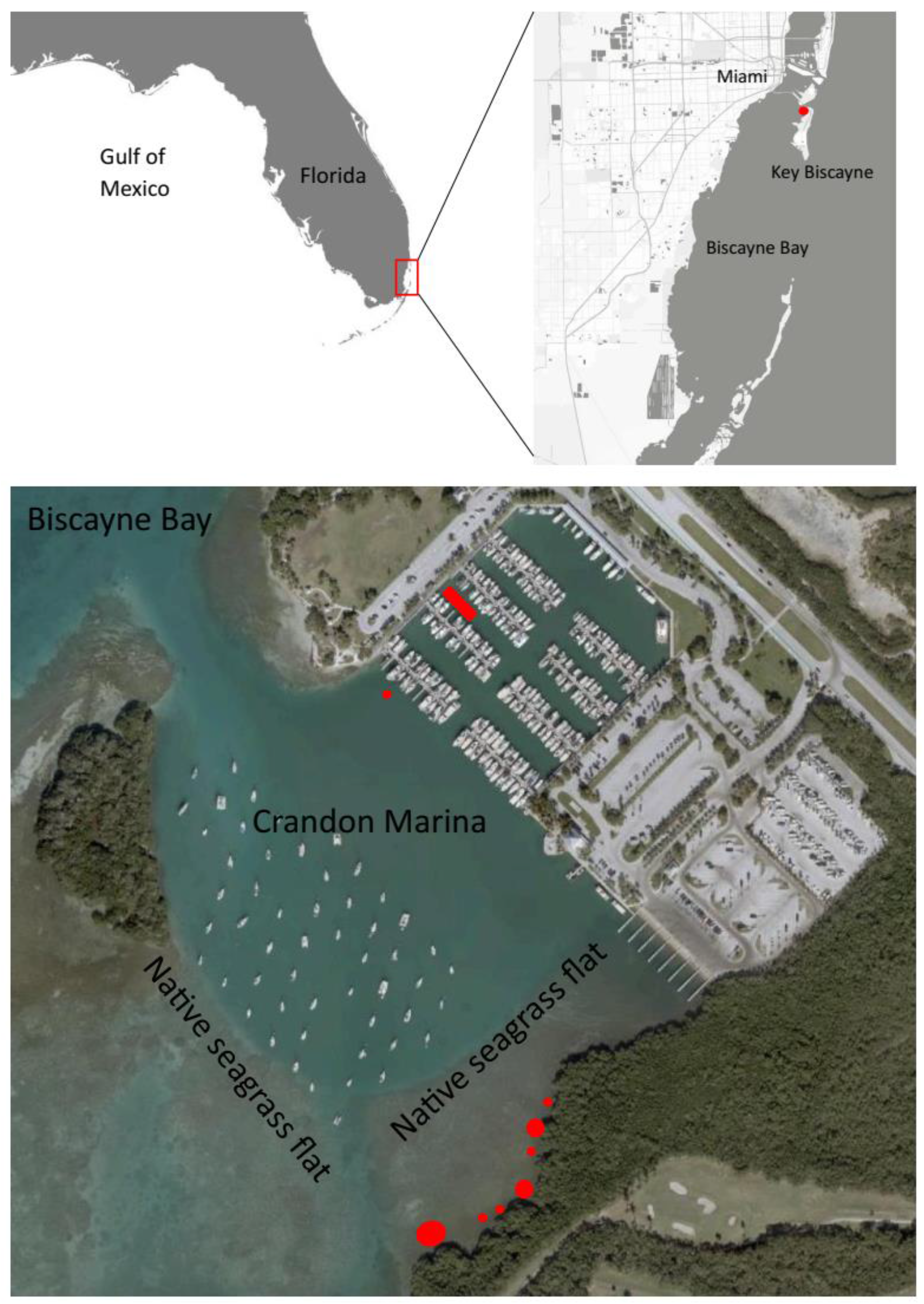
Maps showing the location of Crandon Marina (Key Biscayne, FL). Red shapes indicate the location of *H. stipulacea* (not to scale).

## Results

### Morphological description

Collected seagrass samples displayed paired elongated ellipsoid leaves (2.3 – 5.4cm long,0.5 – 0.8cm wide) with obtuse apices and serrulate margins (Fig. 2). Cross veins were present at approx. 45º angles off the midrib vein and petioles (0.9 – 1.7cm long) were ensheathed in translucent scale leaves. Rhizome thickness was approx. 2mm and root length ranged from 2.8 –10.6 cm. Internode length varied and ranged from 1.1 – 2.0 cm. Flowers were not observed in any of our collected samples.

**Fig 2.**
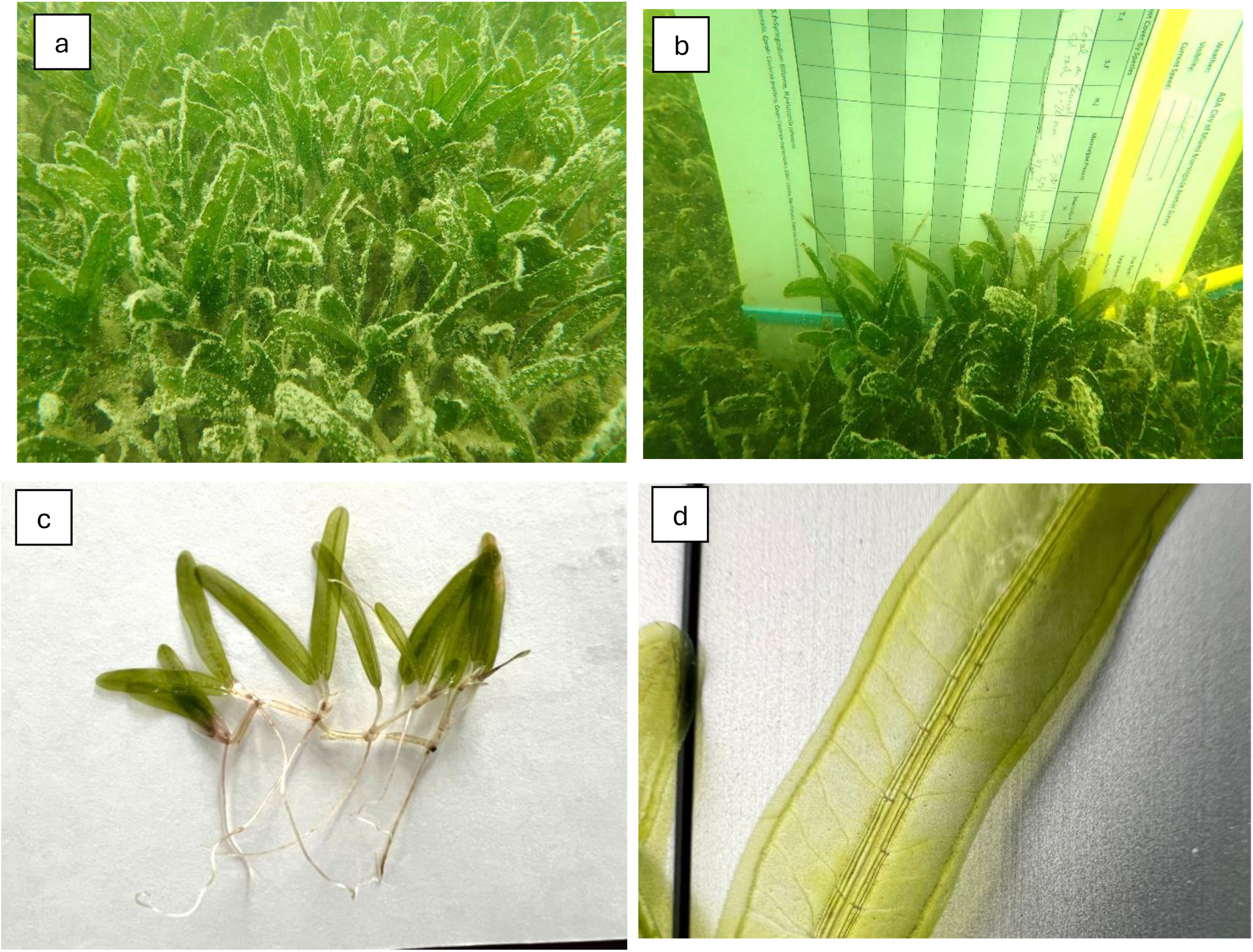
Field photographs of *H. Stipulacea* inside Crandon Marina (Key Biscayne, Florida) (a,b). Close-up detail of samples collected inside the marina, structure and leaf cross veins (c,d). Photo credits: Matthew White (a,b) and Justin Campbell (c,d).

**Fig 3.**
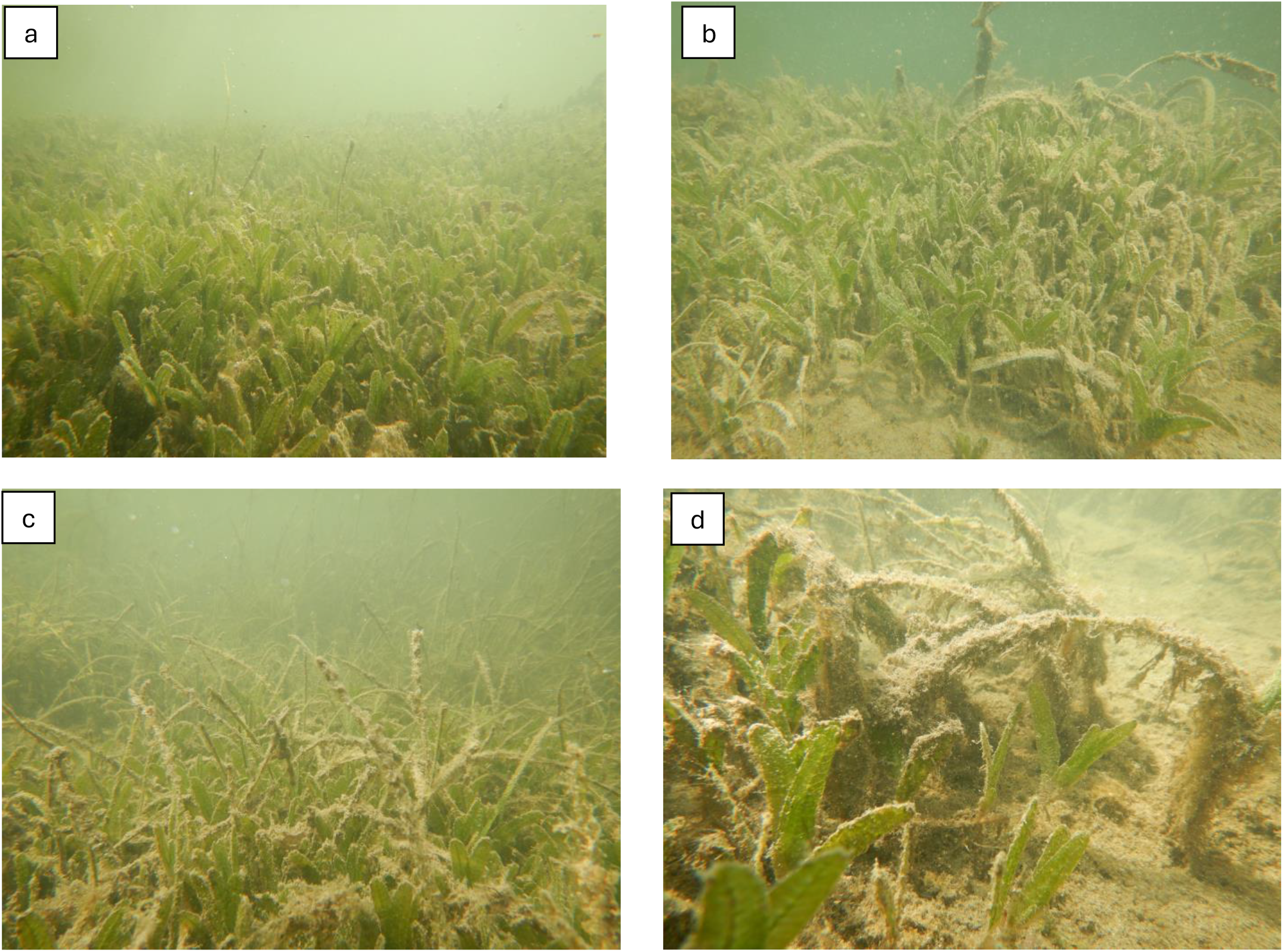
Photographs of *H. Stipulacea* outside of the marina, along the mangrove fringe. Many patches were of mixed composition with the native seagrasses, *T. testudinum, S. filiforme*, and *H. wrightii*. Photo credits, Justin Campbell.

### Survey results

Nine distinct occurrences of *H. stipulacea* were located during our surveys, one large meadow within Crandon Marina and eight smaller patches outside of the marina (see Fig. 1 and Table 1 for characteristics). Inside the marina, the meadow consisted of an irregular mosaic of dense *H. stipulacea* patches, separated by bare silty sediment. No other seagrasses were present; however, some patches did contain several individuals of calcareous green algae (*Halimeda* spp). Averaged across the entire marina meadow, *H. stipulacea* percent cover was 60 ± 12.8% (mean ± SE) and total above-belowground biomass was 360.0 ± 79.2g wet mass / m^2^. Outside of the marina, patches were relatively small (0.8 – 17.4 m^2^), shallow (∼1m depth), and strictly located along the mangrove fringe to the east of the boat anchorage (Fig. 1). Often these meadows were mixed with the native species *Thalassia testudinum, Syringodium filiforme*, and *Halodule wrightii*.

**Table 1.**
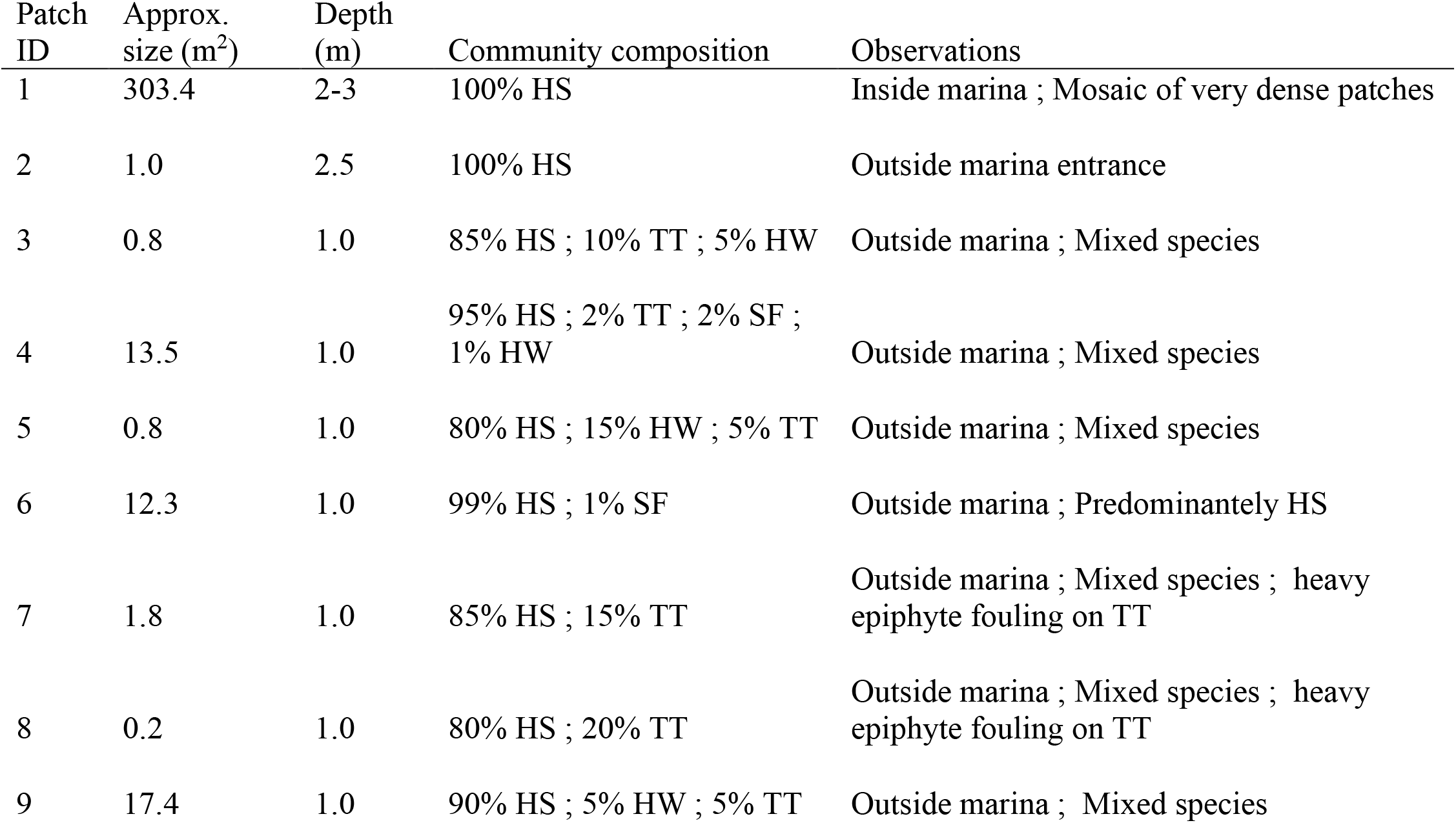
*Halophila stipulacea* characteristics at surveyed sites at Crandon Marina, Key Biscayne, FL. HS= *H. stipulacea*; TT = *T. testudinum*; SF = *S. filiforme*; HW = *H. wrightii*.

## Discussion

Our report is comparable to others from the Caribbean documenting the first sightings of *H. stipulacea* in disturbed habitats such as marinas, ports, or anchorages (Willette et al. 2014).Crandon Marina has ∼250 boats, with the capacity to harbor medium-large sized sailboats (up to 25m in length). These vessels are likely capable of travel to and from areas where *H. stipulacea* is well-established (Puerto Rico, Virgin Islands), thus serving as a potential mechanism for introduction here in south Florida. Our observations are also consistent with others regarding the overall appearance, dense patches approaching 100% substrate cover. The marina meadow was largest and consisted of dense monospecific patches separated by bare substrate. The meadow did not seem to extend towards the deeper central portions of the marina; however, more thorough surveys are required. Outside of the marina, patches were restricted to the shallow sand halo surrounding the mangrove fringe (Fig. 1). While these patches were predominantly comprised of *H. stipulacea*, some were interspersed with lower abundances of native seagrasses (Table 1). It is possible that fragments were transported to the external anchorage by sailboats, and those fragments drifted over the native seagrass flat to the mangrove fringe where they settled in the sand halo and became established. It appears that these patches are now expanding both eastward towards the mangroves and westward toward the flat. The native seagrass flat between the anchorage and the mangrove fringe consists of dense *T. testudinum* and *S. filiforme*, and we did not locate any *H. stipulacea* patches inside this area, only along the mangrove fringe. We assert that it will be important to monitor these fringe patches as they begin to approach the dense flat.

It is difficult to know exactly how long *H. Stipulacea* has been at this marina and the surrounding area. However, if we apply mean lateral patch expansion rates (∼0.5 cm/day along one axis) from other observational studies in the Caribbean (Willette & Ambrose, 2009), we estimate that some of the smaller patches (∼1m diameter) outside the marina may be approximately 200 days old, while the medium patches (∼4m diameter) may be approximately 800 days (2.2 years) old. It is likely that the larger meadow inside the marina has also been there for at least several years. While these calculations don”t necessarily account for non-linear rates of expansion, they may provide a first-order estimate of patch age.

There are also several other large marinas (e.g. Dinner Key Marina, Miami, FL) and anchorages in the Biscayne Bay area that could serve as additional points of introduction, highlighting further concern and warranting expanded surveys. Biscayne Bay harbors extensive seagrass meadows, some of which have been subjected to multiple ecological disturbances (Santos et al. 2020, Santos et al. 2016). Furthermore, the broader Florida Keys National Marine

Sanctuary contains some of the most expansive (∼16,000 km^2^) and protected seagrass meadows in the western hemisphere (Fourqurean & Rutten, 2003). Evaluating the potential impacts of this invasive is of paramount importance. Future work is needed to (1) better understand the current extent of this new introduction, and (2) discern under what environmental conditions *H. stipulacea* can displace native seagrasses (as reviewed in Winters et al. 2020). Will this introduction continue to expand, or will it be restricted to disturbed environments such marinas, ports and anchorages?

## Notes

### Competing Interest Statement

The authors have declared no competing interest.

